# Breast cancer susceptibility: an integrative analysis of genomic data

**DOI:** 10.1101/279984

**Authors:** Simone Mocellin, Sara Valpione, Carlo Riccardo Rossi, Karen Pooley

**Author notes:** Corresponding author: Simone Mocellin Istituto Oncologico Veneto (IOV-IRCCS), Padova, Italy Dept. Surgery Oncology and Gastroenterology, University of Padova, Italy Via Gattamelata 64 35128 Padova, Italy E.

## Abstract

**Background:** Genome wide association studies (GWAS) are greatly accelerating the pace of discovery of germline variants underlying the genetic architecture of sporadic breast cancer predisposition. We have built the first knowledge-base dedicated to this field and used it to generate hypotheses on the molecular pathways involved in disease susceptibility.

**Methods:** We gathered data on the common single nucleotide polymorphisms (SNPs) discovered by breast cancer risk GWAS. Information on SNP functional effect (including data on linkage disequilibrium, expression quantitative trait locus, and SNP relationship with regulatory motifs or promoter/enhancer histone marks) was utilized to select putative breast cancer predisposition genes (BCPGs). Ultimately, BCPGs were subject to pathway (gene set enrichment) analysis and network (protein-protein interaction) analysis.

**Results:** Data from 38 studies (28 original case-control GWAS enrolling 383,260 patients with breast cancer; and 10 GWAS meta-analyses) were retrieved. Overall, 281 SNPs were associated with the risk of breast cancer with a P-value <10E-06 and a minor allele frequency >1%. Based on functional information, we identified 296 putative BCPGs. Primary analysis showed that germline perturbation of classical cancer-related pathways (e.g., apoptosis, cell cycle, signal transduction including estrogen receptor signaling) play a significant role in breast carcinogenesis. Other less established pathways (such as ribosome and peroxisome machineries) were also highlighted. In the main subgroup analysis, we considered the BCPGs encoding transcription factors (n=36), which in turn target 252 genes. Interestingly, pathway and network analysis of these genes yielded results resembling those of primary analyses, suggesting that most of the effect of genetic variation on disease risk hinges upon transcriptional regulons.

**Conclusions:** This knowledge-base, which is freely available and will be annually updated, can inform future studies dedicated to breast cancer molecular epidemiology as well as genetic susceptibility and development.

**Abbreviations:**

GWAS: genome-wide association study
SNP: single nucleotide polymorphism
BCPG: breast cancer predisposition gene
LD: linkage disequilibrium

## Introduction

With a 10-12% lifetime risk, breast cancer is the most common cancer among women with about 1,700,000 new cases and more than 500,000 deaths each year worldwide (1). Breast cancer is a multifactorial disease stemming from a complex interplay between environmental, reproductive/endocrine and genetic risk factors. Dissecting the genetic architecture of breast cancer susceptibility is a pivotal step to understand the cascade of molecular events underlying breast carcinogenesis, which ultimately could lead to better preventive and therapeutic strategies according to the precision medicine principles (2).

Familial aggregation of breast cancer (which occurs in about 10% of cases) has led to family-based linkage analysis and positional cloning studies demonstrating that rare (<1%) germline DNA variation in high to moderate penetrance cancer predisposition genes such as *BRCA1, BRCA2, PTEN, CHEK2, ATM, BRIP1* and *PALB2*-accounts for about 15-20% of the familial risk of this disease (3,4). The residual heritability for breast cancer is believed to be sustained by a polygenic model according to the common disease/common variant hypothesis. Subsequent case-control studies based on the candidate gene approach (also known as hypothesis testing approach) have identified some common germline variants (mainly single nucleotide polymorphisms, SNPs) linked to breast cancer risk, though the evidence quality is often low mainly because of small sample size and result heterogeneity (5,6). More recently, the completion of the Human Genome Project and the implementation of genome-wide association studies (GWAS) – based on a hypothesis generating (also known as data driven) approach and testing hundreds of thousands of SNPs at a time has greatly accelerated the pace of discovery of low penetrance variants linked to the risk of many diseases, including several cancer types (7).

To date, tens of GWAS dedicated to breast cancer have been published, and many single SNPs have been associated with the risk of this malignancy (3,6). This has led to an overwhelming wealth of data which are often difficult to manage by the single reader, in part because most susceptibility loci are intergenic (and thus are linked neither to an obvious gene nor to an obvious functional effect), which hinders a straightforward biological interpretation typical of candidate gene studies.

With the present work we intended to systematically review breast cancer GWAS findings in order to provide readers with the first publicly available knowledge-base dedicated to the relationship between germline genomic DNA variation and breast cancer risk. According to the above mentioned polygenic model of sporadic tumor inheritance and using modern SNP-to-gene and gene-to-function approaches such as integrative analysis of genomic data (8,9) as well as pathway and network analysis (10,11), we also aimed to suggest a biological interpretation of current findings. In particular, we tried to exploit the available GWAS evidence to comprehensively identify the cell pathways whose germline variation condition the predisposition to breast cancer, with an additional effort to prioritize genes/pathways/networks which could be of special relevance to inform future studies in the fields of both molecular epidemiology and biology of breast cancer.

## Materials and Methods

We collected GWAS findings on breast cancer risk (along with other genomic data, see below) to identify breast cancer risk associated SNPs, which were then linked to breast cancer predisposition genes (BCPGs): the data from this knowledge-base were used to perform pathway and network analysis. A flowchart of the study design is reported in **Figure 1**.

**Figure 1:**
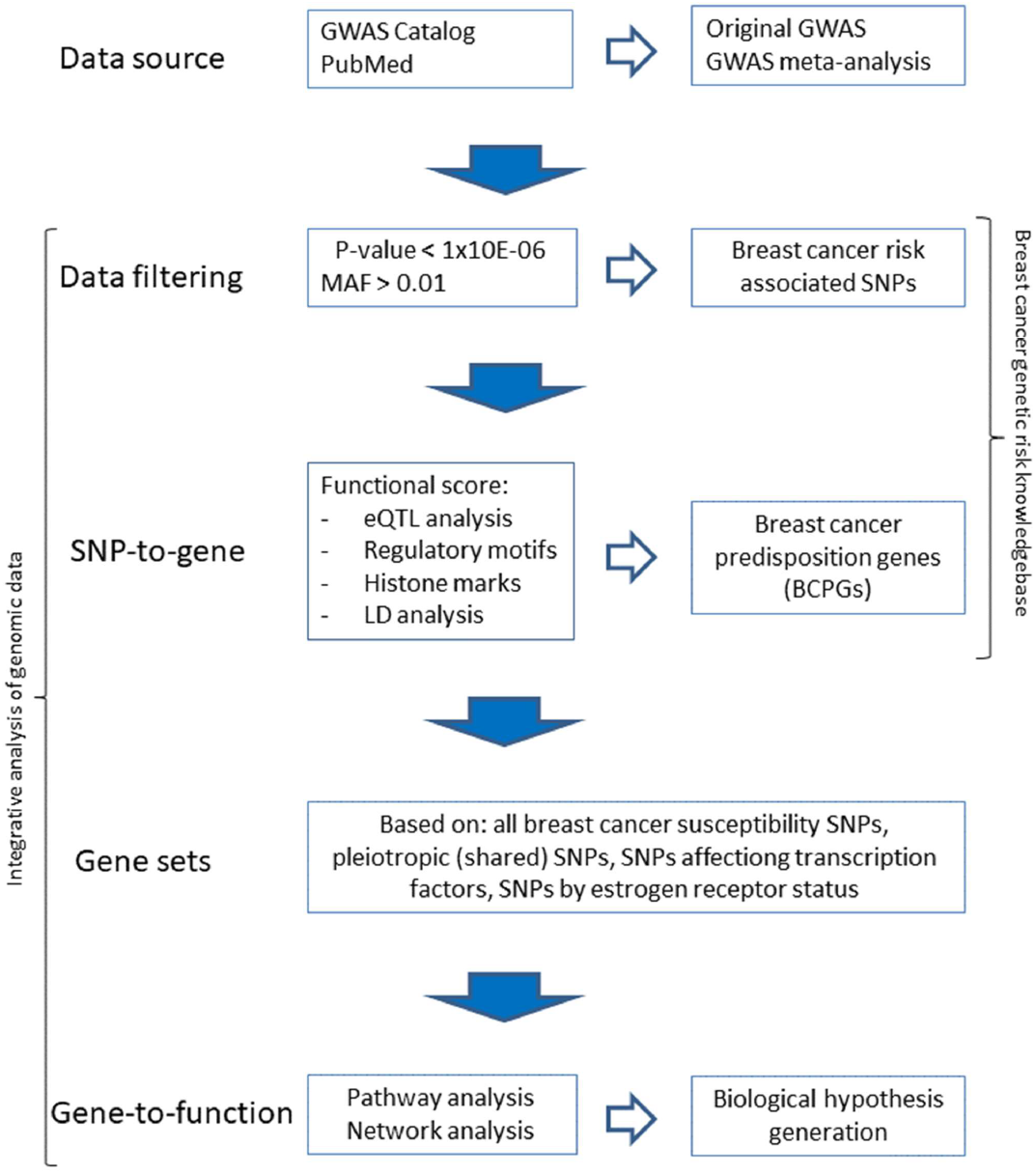
Study design: flow chart of the integrative analysis of genomic data on breast cancer susceptibility

### Breast cancer risk associated SNPs

GWAS addressing the role of germ-line single nucleotide polymorphisms (SNPs) in breast cancer susceptibility were retrieved in the GWAS Catalog repository (7) as well as by performing a systematic review in PubMed (search terms: “breast cancer”, “GWAS”). GWAS meta-analyses were also included for data extraction. Searches were updated until the 11 ?sup>th?/sup> of December 2017.

To be included in the knowledge-base, each SNP had to be associated with breast cancer risk with a nominal P-value lower than 1×10E-06 (genome-wide significance level) and have a minor allele frequency ≥1% in the general population of European ancestry.

### SNP-to-gene analysis

Following the principles of integrative analysis of genomic data (8,9), the functional association between a breast cancer risk associated SNP and a gene (hereafter called BCPG) was scored according to three types of information:

A. SNP relationship with gene(s): [**Category 1 – score=2**] This applies to within-gene non-synonymous variants (e.g., missense SNPs), variants associated with expression quantitative trait locus (eQTL) data (based on GTex portal database (12)), variants in high linkage disequilibrium (LD) (pairwise r-squared ≥ 0.8) with another SNP that is an eQTL hit, or variants in high LD with a within-gene non-synonymous variant; [**Category 2 – score=1**] This applies to within-gene synonymous variants, or variants located in a non coding gene region (e.g., intronic SNPs), or variants in high LD with another within gene SNP (synonymous variant, or variant located in a non-coding gene region); [**Category 3 – score=0**] intergenic and non eQTL hit variants.
B. SNP relationship with regulatory motifs (i.e., variant potentially affecting the binding of transcription factors based on a library of position weight matrices scored on genomic sequences (13)): [**Category 1 – score=1**] variant changing these motifs, or variant in high LD with another SNP changing these motifs; [**Category 2-score = 0**] no change of motif/LD with SNP changing motif.
C. SNP relationship with breast tissue specific promoter/enhancer histone marks (based on data from the Roadmap Epigenomics Project (14)): [**Category 1, score=1**] variant co-localization with these marks, or variant in high LD with another SNP co-localising with these marks; [**Category 2, score=0**] no co-localization with epigenetic marks/LD with SNP colocalising with epigenetic marks.

The principles underlying the above scoring system are analogous to those employed in well-established functional annotation databases (such as RegulomeDB (15) and HaploReg (16)). However, we added the information deriving from LD analysis (which was performed using the LDLink website (17)), which increases the likelihood of identifying additional functional variants relevant to disease susceptibility.

### Gene-to-function: pathway and network analysis

Once BCPGs were identified, we used them to perform pathway and network analysis in order to identify biological functions whose genetic perturbations can predispose to breast cancer development (10,11).

For pathway analysis purposes we utilized gene set enrichment analysis (GSEA) as performed by the EnrichR web server (18). Hypergeometric distribution was used to calculate the statistical significance of gene overlapping (19), followed by correction for multiple hypotheses testing (using the false discovery rate [FDR] method (20)). Only pathways with a FDR <0.05 were considered of interest.

Also protein-protein interaction (PPI) networks can be employed to select gene sets. In contrast to pathways, networks are not based on specific biological functions but are built on the basis of both direct (physical) and indirect (genetic) interactions between gene products (proteins). For network analysis, we utilized the STRING 10.5 web server (21). In order to consider only highly reliable information on protein-protein interactions (PPI), we set the interaction score to ≥ 0.7 (high confidence). The resulting network provides information of the degree of overall connectivity across imputed gene products (as quantified by the ratio between observed and expected interactions [a.k.a“edges”] between proteins [a.k.a.“nodes”], and formally tested by means of a PPI enrichment test). Then, molecular clusters (subnetworks) can be identified that can be utilized for gene set enrichment analysis (only subnetworks with at least three BCPGs were considered). When the network connectivity is low, the PPI database can be exploited to add first-shell interactors (we chose to add no more than 10 such interactors to avoid data over-interpretation) and then re-run pathway analysis. Ultimately, this data augmentation process increases the likelihood of identifying relevant biological pathways which would be otherwise overlooked when starting with only few BCPGs belonging to a given pathway.

### Other analyses

Within the frame of network analysis, we searched for so called “hub proteins”: these are molecules with a high degree of connectivity with the other network components and thus are likely to play a dominant role in the activity of the network itself (they are also known as “influencers”) (22). To this aim, we used the EsyN webtool (23) to calculate the collective influence score, which defined as the product of a node-reduced degree (number of edges minus one) times the sum of the reduced degree of the nodes that are two steps away (a.k.a. radius) (24).

Finally, in order to provide further information beyond the cis effects of included variants (as done in the above analyses), we explored the potential effect *in trans* of breast cancer associated SNPs. To this aim, we first identified the transcription factors among the putative BCPGs: then, the genes whose expression were regulated by these transcription factors (identified by using the Uniprot (25) and TRRUST (26) repositories) were input in both pathway and network analysis to assess the cellular functions potentially affected by germline variation linked to cancer risk.

## Results

### Breast cancer risk associated SNPs

We found 38 studies (published between 2007 and 2017) which matched our inclusion criteria (27–64). Of these, 28 were original case-control GWAS (overall enrolling 383,260 patients with breast cancer) and 10 were meta-analyses of previously published case-control GWAS (overall enrolling 239,271 patients with breast cancer) (**Supplementary Table 1**).

In most studies, patients and controls were of European ancestry (71% among original GWAS; 97% among GWAS meta-analyses); in the remaining studies, individuals were mainly of Asian ancestry among original GWAS (96%) and African-American among GWAS meta-analyses (100%). Only one original study was dedicated to male breast cancer. As regards tumor subtype by estrogen receptor expression, two original GWAS were dedicated to receptor negative and two to receptor positive breast cancer, whereas four GWAS meta-analyses were dedicated to ER negative cases. In the original articles associations were reported (and are reported hereafter in the text) as per-allele odds ratios (ORs).

Overall, 281 SNPs were associated with the risk of breast cancer with a P-value <and a MAF >0.01 (**Supplementary Table 2**); the median minor allele frequency was 0.28 (interquartile range [IQR]: 0.16-0.39); the median OR was 0.93 (IQR: 0.91-0.95) and 1.11 (IQR: 1.06-1.19) for protective and risk alleles, respectively.

Chromosome distribution showed an over-representation of chromosome 5 (signals observed: 32; expected: 15; FDR: 0.0003) and chromosome 19 (signals observed: 12; expected: 5; FDR: 0.014).

Linkage disequilibrium (LD) analysis of the 281 SNPs showed that 48 polymorphisms were tagged by one or more other variants (LD r-squared >0.8), leading to the identification of 233 breast cancer predisposition loci (**Supplementary Table 3**).

Out of 281 reported SNPs associated with breast cancer risk at a genome-wide significance level, only 34 (12.1%) were reported by two or more data sources.

Whereas most studies (n=21, 55.3%) enrolled women with unspecified sporadic breast cancer, subgroups were specifically investigated by others: estrogen receptor negative breast cancer (n=7); estrogen receptor positive tumor (n=2); triple negative tumor (n=1); breast cancer in BRCA1/2 mutation carriers (studies, n=3); early onset breast cancer (n=1); breast cancer in post-menopausal women (n=1); lobular carcinoma (n=1); and breast cancer in males (n=1).

### Breast cancer predisposition genes

The majority of SNPs were located within coding genes (n=160, 56.9%). More specifically, SNPs were located in gene 3’-UTR (n=7, 2.5%), intron (n=140, 49.8%), exon (n=13, 4.6%; of these: missense, n=8, synonymous, n=4 and non-sense [stop gain], n=1). The remaining SNPs were intergenic region (n=95, 33.8%) and within non-coding genes (n=17, 9.2%). Of note, 6 intergenic SNPs (2.1%) were in high LD with non-tested SNPs located within a gene and other 8 SNPs (2.8%) were in high LD with non-tested missense variants.

As regards eQTL analysis, 107 SNPs (38.1%) were directly associated with a significant effect on the expression of one or more genes, and 3 SNPs (1.1%) were in high LD with SNPs with an eQTL effect. A large majority of 229 variants (81.5%) were associated with changing regulatory motifs, with 43 SNPs (15.3%) in high LD with those 229 variants, whereas only 9 SNPs had no impact on regulatory motifs. In addition, 79 SNPs (28.1%) co-localized with promoter/enhancer histone marks, with 107 variants (38.1%) in high LD with those 79 SNPs, and 95 SNPs (33.8%) having no such property.

Based on the above information, we associated the 281 SNPs linked with breast cancer risk to 334 genes with a score ranging from 0 to 4 (**Supplementary Table 4**): SNPs with low (0-1), intermediate (2) and high (3-4) functional score were 30 (10.7%), 68 (24.2%), and 183 (65.1%), respectively. In order to exclude genes with low level of evidence of association with breast cancer risk SNPs, we further considered only SNPs with a score equal or greater than 2 (n=251). With this cut off, we identified 296 putative BCPGs, which were the genes utilized in the following primary analysis. These genes code for known proteins in most cases (n=255,86.1%).

### Pathway and network analysis

Primary gene set enrichment analysis demonstrated that the 296 BCPGs are enriched in genes involved in apoptotic pathway and peroxisome machinery, as illustrated in **Table 1**.

**Table 1:**
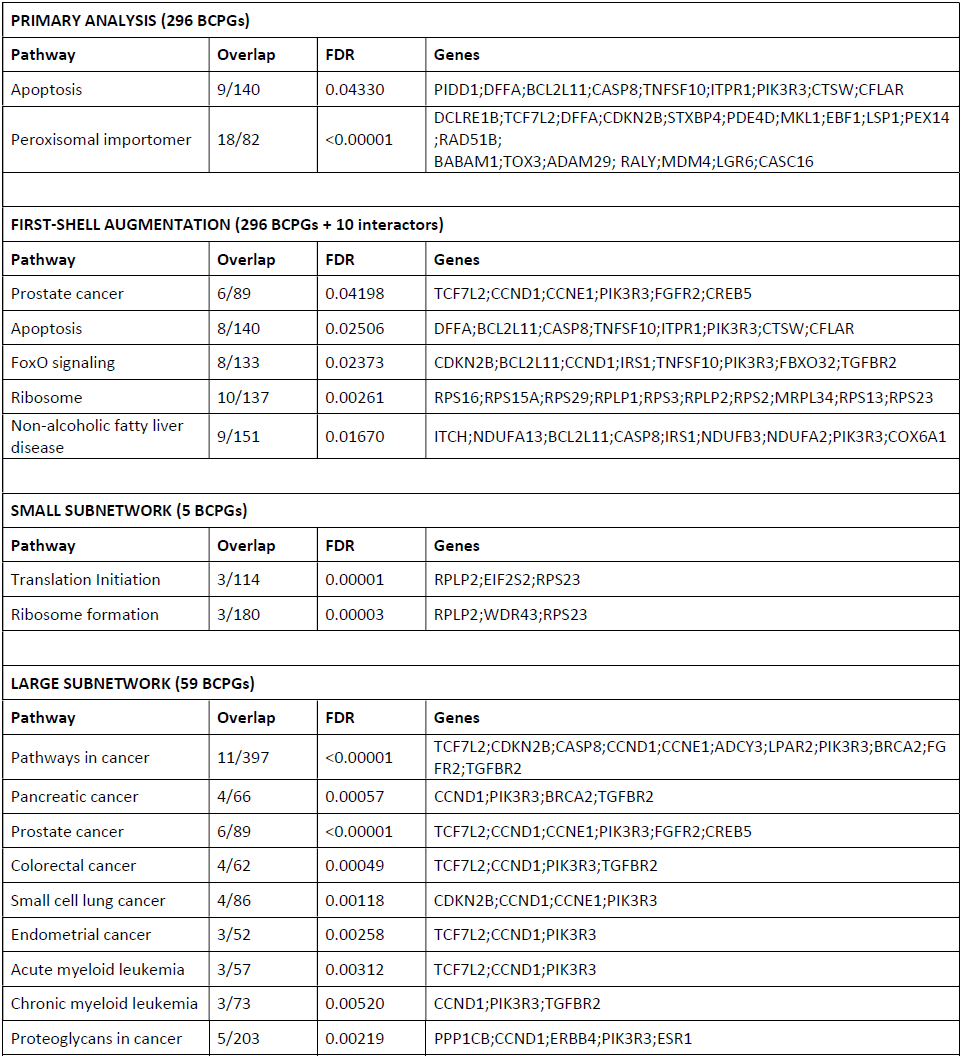
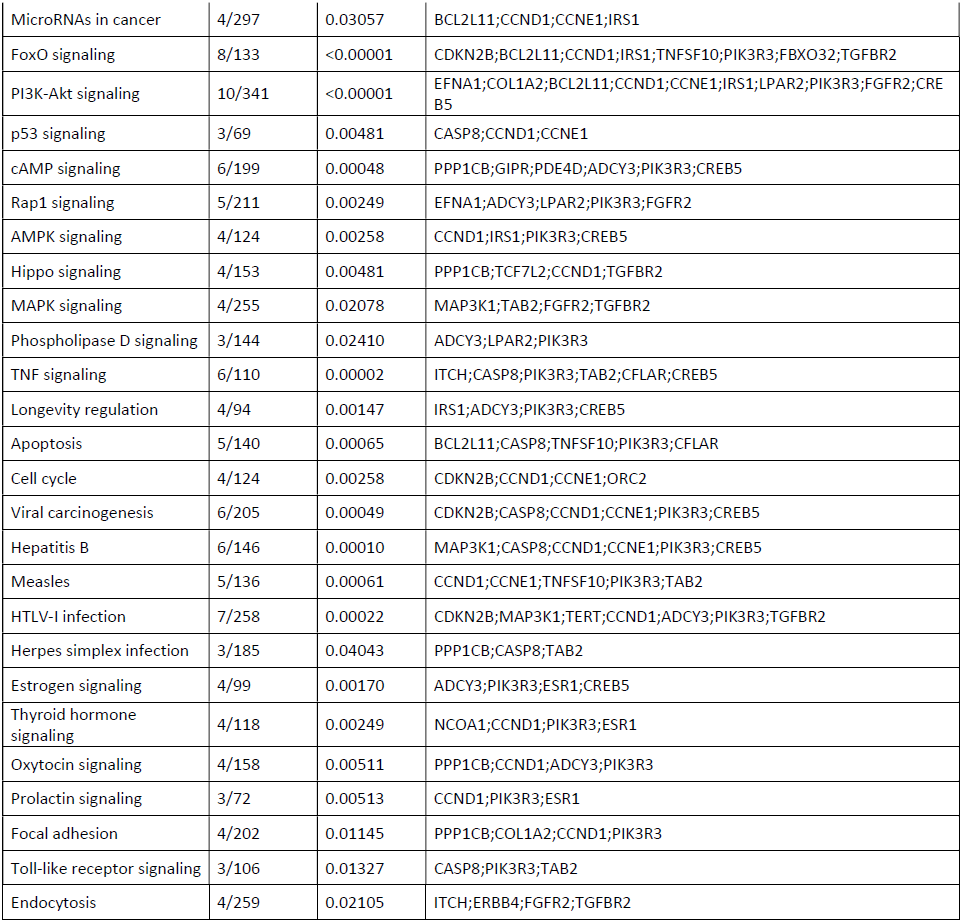
Pathway analysis of breast cancer predisposition genes (BCPGs). Overlap: number of BCPGs over number of pathway genes. FDR: false discovery rate.

| PRIMARY ANALYSIS (296 BCPGs) |  |  |  |
| --- | --- | --- | --- |
| Pathway | Overlap | FDR | Genes |
| Apoptosis | 9/140 | 0.04330 | PIDD1;DFFA;BCL2L11;CASP8;TNFSF10;ITPR1;PIK3R3;CTSW;CFLAR |
| Peroxisomal importomer | 18/82 | <0.00001 | DCLRE1B;TCF7L2;DFFA;CDKN2B;STXBP4;PDE4D;MKL1;EBF1;LSP1;PEX14;RAD51B;BABAM1;TOX3;ADAM29; RALY;MDM4;LGR6;CASC16 |
| FIRST-SHELL AUGMENTATION (296 BCPGs + 10 interactors) |  |  |  |
| Pathway | Overlap | FDR | Genes |
| Prostate cancer | 6/89 | 0.04198 | TCF7L2;CCND1;CCNE1;PIK3R3;FGFR2;CREB5 |
| Apoptosis | 8/140 | 0.02506 | DFFA;BCL2L11;CASP8;TNFSF10;ITPR1;PIK3R3;CTSW;CFLAR |
| FoxO signaling | 8/133 | 0.02373 | CDKN2B;BCL2L11;CCND1;IRS1;TNFSF10;PIK3R3;FBXO32;TGFB2 |
| Ribosome | 10/137 | 0.00261 | RPS16;RPS15A;RPS29;RPLP1;RPS3;RPLP2;RPS2;MRPL34;RPS13;RPS23 |
| Non-alcoholic fatty liver disease | 9/151 | 0.01670 | ITCH;NDUFA13;BCL2L11;CASP8;IRS1;NDUFB3;NDUFA2;PIK3R3;COX6A1 |
| SMALL SUBNETWORK (5 BCPGs) |  |  |  |
| Pathway | Overlap | FDR | Genes |
| Translation Initiation | 3/114 | 0.00001 | RPLP2;EIF2S2;RPS23 |
| Ribosome formation | 3/180 | 0.00003 | RPLP2;WDR43;RPS23 |
| LARGE SUBNETWORK (59 BCPGs) |  |  |  |
| Pathway | Overlap | FDR | Genes |
| Pathways in cancer | 11/397 | <0.00001 | TCF7L2;CDKN2B;CASP8;CCND1;CCNE1;ADCY3;LPAR2;PIK3R3;BRCA2;FGFR2;TGFB2 |
| Pancreatic cancer | 4/66 | 0.00057 | CCND1;PIK3R3;BRCA2;TGFB2 |
| Prostate cancer | 6/89 | <0.00001 | TCF7L2;CCND1;CCNE1;PIK3R3;FGFR2;CREB5 |
| Colorectal cancer | 4/62 | 0.00049 | TCF7L2;CCND1;PIK3R3;TGFB2 |
| Small cell lung cancer | 4/86 | 0.00118 | CDKN2B;CCND1;CCNE1;PIK3R3 |
| Endometrial cancer | 3/52 | 0.00258 | TCF7L2;CCND1;PIK3R3 |
| Acute myeloid leukemia | 3/57 | 0.00312 | TCF7L2;CCND1;PIK3R3 |
| Chronic myeloid leukemia | 3/73 | 0.00520 | CCND1;PIK3R3;TGFB2 |
| Proteoglycans in cancer | 5/203 | 0.00219 | PPP1CB;CCND1;ERBB4;PIK3R3;ESR1 |
| MicroRNAs in cancer | 4/297 | 0.03057 | BCL2L11;CCND1;CCNE1;IRS1 |
| FoxO signaling | 8/133 | <0.00001 | CDKN2B;BCL2L11;CCND1;IRS1;TNFSF10;PIK3R3;FBXO32;TGFB2 |
| PI3K-Akt signaling | 10/341 | <0.00001 | EFNA1;COL1A2;BCL2L11;CCND1;CCNE1;IRS1;LPAR2;PIK3R3;FGFR2;CREB5 |
| p53 signaling | 3/69 | 0.00481 | CASP8;CCND1;CCNE1 |
| cAMP signaling | 6/199 | 0.00048 | PPP1CB;GIPR;PDE4D;ADCY3;PIK3R3;CREB5 |
| Rap1 signaling | 5/211 | 0.00249 | EFNA1;ADCY3;LPAR2;PIK3R3;FGFR2 |
| AMPK signaling | 4/124 | 0.00258 | CCND1;IRS1;PIK3R3;CREB5 |
| Hippo signaling | 4/153 | 0.00481 | PPP1CB;TCF7L2;CCND1;TGFB2 |
| MAPK signaling | 4/255 | 0.02078 | MAP3K1;TAB2;FGFR2;TGFB2 |
| Phospholipase D signaling | 3/144 | 0.02410 | ADCY3;LPAR2;PIK3R3 |
| TNF signaling | 6/110 | 0.00002 | ITCH;CASP8;PIK3R3;TAB2;CFLAR;CREB5 |
| Longevity regulation | 4/94 | 0.00147 | IRS1;ADCY3;PIK3R3;CREB5 |
| Apoptosis | 5/140 | 0.00065 | BCL2L11;CASP8;TNFSF10;PIK3R3;CFLAR |
| Cell cycle | 4/124 | 0.00258 | CDKN2B;CCND1;CCNE1;ORC2 |
| Viral carcinogenesis | 6/205 | 0.00049 | CDKN2B;CASP8;CCND1;CCNE1;PIK3R3;CREB5 |
| Hepatitis B | 6/146 | 0.00010 | MAP3K1;CASP8;CCND1;CCNE1;PIK3R3;CREB5 |
| Measles | 5/136 | 0.00061 | CCND1;CCNE1;TNFSF10;PIK3R3;TAB2 |
| HTLV-I infection | 7/258 | 0.00022 | CDKN2B;MAP3K1;TERT;CCND1;ADCY3;PIK3R3;TGFB2 |
| Herpes simplex infection | 3/185 | 0.04043 | PPP1CB;CASP8;TAB2 |
| Estrogen signaling | 4/99 | 0.00170 | ADCY3;PIK3R3;ESR1;CREB5 |
| Thyroid hormone signaling | 4/118 | 0.00249 | NCOA1;CCND1;PIK3R3;ESR1 |
| Oxytocin signaling | 4/158 | 0.00511 | PPP1CB;CCND1;ADCY3;PIK3R3 |
| Prolactin signaling | 3/72 | 0.00513 | CCND1;PIK3R3;ESR1 |
| Focal adhesion | 4/202 | 0.01145 | PPP1CB;COL1A2;CCND1;PIK3R3 |
| Toll-like receptor signaling | 3/106 | 0.01327 | CASP8;PIK3R3;TAB2 |
| Endocytosis | 4/259 | 0.02105 | ITCH;ERBB4;FGFR2;TGFB2 |

Network analysis suggested that BCPGs protein products did not have more interactions among themselves than expected for a random protein set of equal size drawn from the proteome (observed edges: 98; expected edges: 83; PPI enrichment test P-value: 0.0527), indicating that these proteins are not remarkably biologically connected as a group. When 10 first-shell interactors were added to the network, ribosome proteins were then included in the enrichment list (**Table 1**).

Network analysis also enabled us to identify one large (n=59) and one small (n=5) subnetwork (**Figure 2**): the former was enriched in several cancer-related pathways, including apoptosis and estrogen receptor signaling, whereas the latter was enriched in ribosome machinery components (see **Table 1**). Finally, influence analysis of the large subnetwork identified estrogen receptor 1 (ESR1) as the most influential protein (**Suppementary Table 5**).

**Figure 2:**
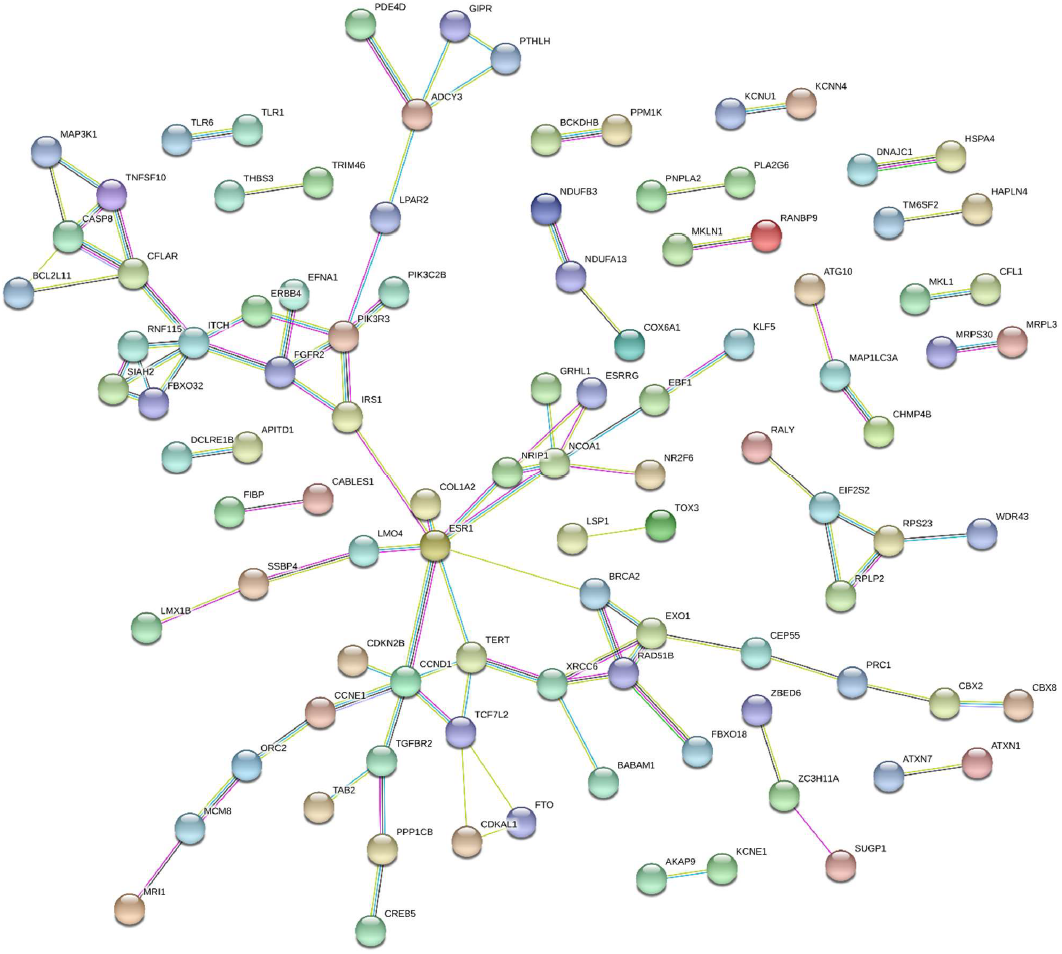
Network analysis: network plot of protein-protein interactions regarding the products of the putative breast cancer predisposition genes identified through the integrative analysis of GWAS data

### Subgroup analysis

In a first subgroup analysis we considered the BCPGs encoding transcription factors: there were 36 such genes (**Figure 3**), which represent 12.2% of the BCPGs identified in this work, a figure only slightly higher than expected (10%). These transcription factors target 252 genes (**Figure 3**), with nine also being BCPGs (*AHRR, BRCA2, CCND1, CDKN2B, ESR1, FOXP1, FTO, LPAR2, TERT*).

**Figure 3:**
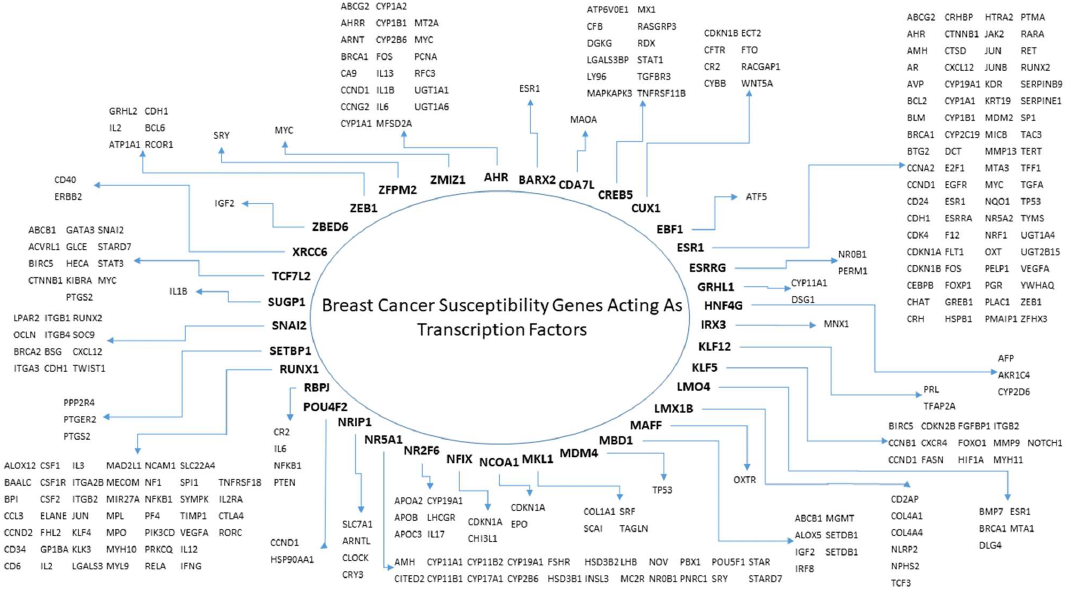
Breast cancer susceptibility regulons: targets of transcription factors whose germline variation is associated with breast cancer risk

Pathway analysis of the 252 targets demonstrated a significant enrichment in many cancer-related pathways, including those involved in the pathogenesis of different tumor types (mainly but not only carcinomas), cell cycle and apoptosis, multiple signaling pathways (such as p53, PI3K-Akt, Wnt, Hippo, Mapk, ErbB, HIF-1 and VEGF), hormone pathways (including sex hormones), immunity (with special regard to anti-viral immune response), and cell adhesion (**Table 2**).

**Table 2:**
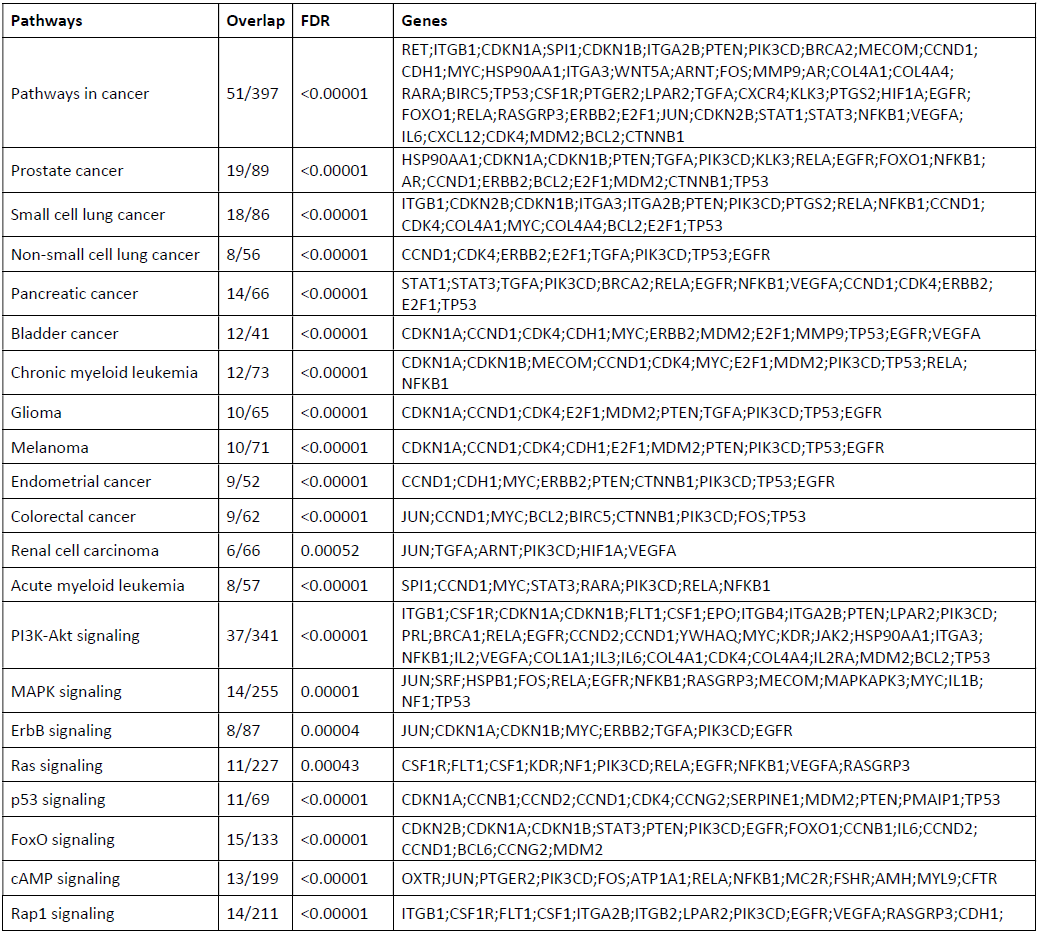
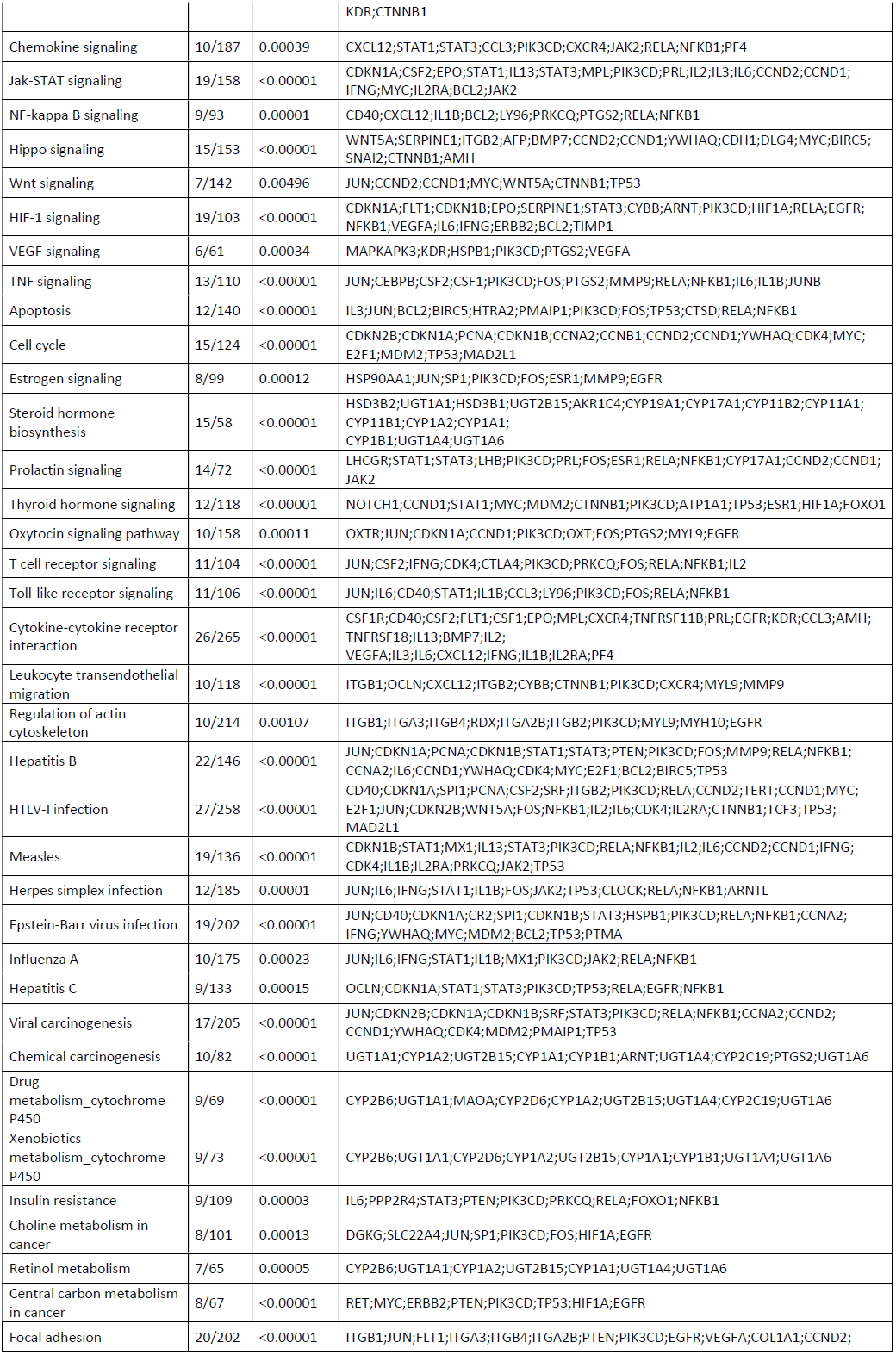
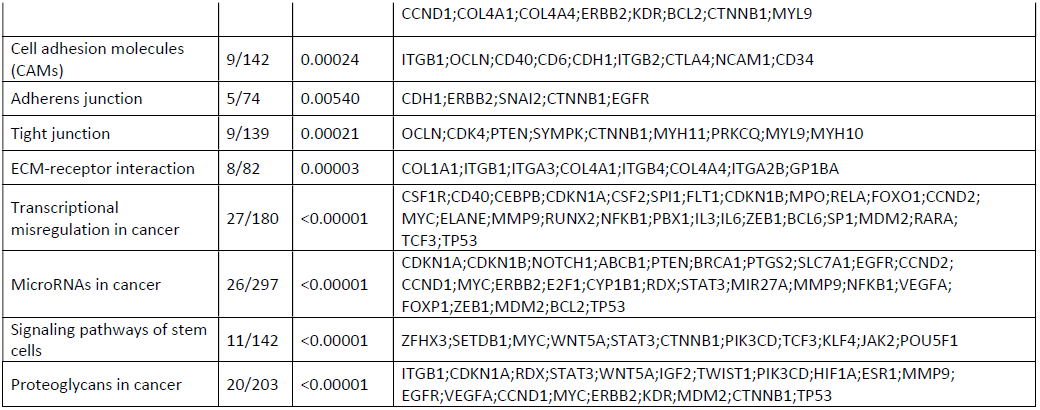
Pathway analysis of breast cancer predisposition genes (BCPGs) encoding transcription factors. Overlap: number of BCPGs over number of pathway genes. FDR: false discovery rate.

Network analysis revealed a very high degree of connectivity across these target genes (observed edges: 3105; expected: 1104; PPI enrichment p-value: <10E-16); influence analysis showed that the top ten most influential proteins largely overlapped with those identified in the primary analysis (8/10), with ESR1 being the second ranking molecule (**Suppementary Table 5**).

Data were available for 238 SNPs linked to 275 genes which also allowed us to perform a subgroup analysis dedicated to estrogen receptor negative breast cancer (only four SNPs were available for estrogen receptor positive cases). Pathway and network analysis yielded results very similar to those obtained in the primary analysis where all breast cancer cases (both receptor positive and negative) were included (data not shown), likely because of the high degree of overlapping between the SNPs (and consequently of genes) of the two series.

### SNPs shared with other tumors

Finally, we assessed whether some breast cancer risk associated SNPs are shared with other malignancies, a phenomenon known as pleiotropy (65). Querying the GWAS Catalog, we found 37 breast cancer risk SNPs shared with other eight tumor types (details are reported in **Table 3**): ovarian carcinoma (n=7), prostate carcinoma (n=4), lung carcinoma (n=2), thyroid carcinoma (n=1), esophageal carcinoma (n=1), renal cell carcinoma (n=1), cutaneous melanoma (n=1), glioma/glioblastoma (n=1) and a tumor miscellany mainly including ovarian, prostate and lung carcinoma (n=28). In two cases, the breast cancer susceptibility locus was shared with other three tumor types: one SNP (rs13016963) was located in chromosome 2q33.1 (sharing tumors: prostate and esophageal carcinomas, and cutaneous melanoma), the other SNP (rs10069690) in chromosome 5p15.33 (sharing tumors: ovarian and thyroid carcinomas, and glioma/glioblastoma).

**Table 3:** Breast cancer risk associated single nucleotide polymorphisms (SNPs) shared with other malignancies

| Cancer | Chromosome | SNP | Genes |
| --- | --- | --- | --- |
| Glioma/Glioblastoma | 5p15.33 | rs10069690 | TERT |
| Ovarian carcinoma | 5p15.33 | rs10069690 | TERT |
| Thyroid carcinoma | 5p15.33 | rs10069690 | TERT |
| Miscellany | 9p21.3 | rs1011970 | CDKN2B |
| Prostate carcinoma | 9p21.3 | rs1011970 | CDKN2B |
| Miscellany | 19p13.11 | rs10419397 | ABHD8;ANKLE1;BABAM1 |
| Ovarian carcinoma | 19p13.11 | rs10419397 | ABHD8;ANKLE1;BABAM1 |
| Miscellany | 14q24.1 | rs10483813 | RAD51B |
| Miscellany | 10q26.13 | rs1078806 | FGFR2 |
| Miscellany | 10q26.13 | rs11200014 | FGFR2 |
| Lung carcinoma | 13q13.1 | rs11571833 | BRCA2 |
| Miscellany | 14q24.1 | rs11844632 | RAD51B |
| Miscellany | 20q11.22 | rs11907546 | CHMP4B |
| Miscellany | 10q26.13 | rs1219648 | FGFR2 |
| Esophageal carcinoma | 2q33.1 | rs13016963 | CASP8;ALS2CR12 |
| Melanoma (cutaneous) | 2q33.1 | rs13016963 | CASP8;ALS2CR12 |
| Prostate carcinoma | 2q33.1 | rs13016963 | CASP8;ALS2CR12 |
| Miscellany | 3p24.1 | rs1352941 | NEK10;SLC4A7 |
| Miscellany | 19p13 | rs1469713 | MAU2;SUGP1;NDUFA13;GATAD2A;CILP2;TM6SF2 |
| Prostate carcinoma | 5p15.33 | rs2242652 | TERT |
| Ovarian carcinoma | 19p13.11 | rs2363956 | ABHD8;ANKLE1;MRPL34;OCEL1 |
| Miscellany | 3p24.1 | rs2590265 | NEK10 |
| Miscellany | 10q26.13 | rs2912780 | FGFR2 |
| Miscellany | 10q26.13 | rs2981575 | FGFR2 |
| Miscellany | 10q26.13 | rs2981582 | FGFR2 |
| Miscellany | 10q26.13 | rs3135718 | FGFR2 |
| Prostate carcinoma | 1q32.1 | rs4245739 | MDM4;PIK3C2B |
| Miscellany | 19p13.11 | rs4808075 | ANO8;ABHD8;ANKLE1;BABAM1;OCEL1 |
| Ovarian carcinoma | 19p13.11 | rs4808075 | ANO8;ABHD8;ANKLE1;BABAM1;OCEL1 |
| Miscellany | 3p24.1 | rs481519 | NEK10 |
| Miscellany | 3p24.1 | rs571978 | NEK10 |
| Miscellany | 3p24.1 | rs580057 | NEK10 |
| Miscellany | 5q11.2 | rs59957907 | C5orf67;MAP3K1 |
| Miscellany | 19p13.11 | rs61494113 | ABHD8;ANKLE1;OCEL1 |
| Ovarian carcinoma | 19p13.11 | rs61494113 | ABHD8;ANKLE1;OCEL1 |
| Miscellany | 14q24.1 | rs61986943 | RAD51B |
| Ovarian carcinoma | 9p34 | rs635634 | SURF6;ABO |
| Miscellany | 5q11.2 | rs6450401 | MAP3K1;SETD9 |
| Miscellany | 5q11.2 | rs6890270 | MIER3;SETD9 |
| Miscellany | 14q24.1 | rs71423318 | RAD51B |
| Renal cell carcinoma | 11q22.3 | rs74911261 | KDELC2 |
| Miscellany | 5q11.2 | rs7709971 | C5orf67;MAP3K1 |
| Miscellany | 5q11.2 | rs7714232 | C5orf67;MAP3K1 |
| Lung carcinoma | 5p15.33 | rs7726159 | TERT |
| Miscellany | 5p15.33 | rs7726159 | TERT |
| Ovarian carcinoma | 19p13.11 | rs8170 | PLVAP;NR2F6;BABAM1;MRPL34;USHBP1;ABHD8;ANKLE1 |

These shared SNPs were associated with 34 genes: when we input these BCPGs into a pathway analysis, enrichment in apoptosis and cancer-related pathways was observed (**Supplementary Table 6**). Upon network analysis, the connectivity was very low (observed edges: 2; expected edges: 1; PPI enrichment P-value:24 0.386). Adding 10 first-shell interactors showed the enrichment in cancer-related pathways as well as ribosome machinery and degenerative disease pathways (**Supplementary Table 6**).

## Discussion

We reported on the first knowledge-base dedicated on GWAS-based evidence linking common germline variants to the risk of breast cancer. The information on breast cancer risk associated SNPs forms a knowledge-base which will be made publicly available at our cancer dedicated website (www.mmmp.org(66)) and will be annually updated.

Following the principles of integrative analysis of genomic data, we combined genome-wide information from different sources (e.g., high-throughput genotyping experiments, eQTL analysis, LD analysis, and so on) to make the most of the available evidence (8,9). This is of particular relevance because most SNPs do not have a direct functional effect, indeed a large proportion of associated SNPs are not in the coding regions of genes, and thus additional information is needed to link them to a gene. Then, we used these data to make tentative inferences on the pathways (and most influential molecules within them) whose variation can affect the risk of developing breast cancer.

Data from almost 400,000 women affected with breast cancer showed that 281 SNPs are significantly associated with the risk of this disease, which reduced to 233 risk loci when linkage disequilibrium was taken into account. These findings add new information to the already existing recent literature reviews on this subject, which report up to 172 common variants linked to breast cancer susceptibility (3,4,6,67–71). These SNPs are estimated to account 15-20% of the genetic component of disease risk (3,72), which clearly implies that much more work is needed to fully elucidate the molecular basis of breast cancer predisposition. It has been argued that future GWAS will not lead to the discovery of many more risk variants (3). This appears especially true in terms of rare variants (that is, variants with a MAF <1%) (3,72), as GWAS studies are designed to identify only common polymorphisms (MAF >1%) through a tagging strategy (tested tag-SNPs are in high linkage disequilibrium with non-tested SNPs). Moving forward, massively parallel sequencing technology (a.k.a. next generation sequencing [NGS], which can directly interrogate every genomic position) could provide investigators with the right tool to overcome the challenging hurdle of interrogating rarer variants which may affect risk, thus adding essential information to this field of investigation (73).

The data collected in our knowledge-base can be used to build polygenic predictive models and thus help optimize breast cancer secondary prevention programs (i.e. early detection by mammographic screening) by selecting women at higher risk (74–77). So far, such models have yielded generally unsatisfactory results, as their accuracy remains too low to be clinically implemented. This could be due to the fact that the complex genetic architecture of sporadic breast cancer predisposition remains still to be fully elucidated, as well as to the lack of information on gene-environment interactions (78,79). Nevertheless, the systematic collection of variants associated with breast cancer risk, along with information on their functional effect (as proposed in our knowledge-base) is the first step to build more effective predictive tools.

We utilized the collected information to generate tentative mechanistic hypotheses on the pathways whose perturbation (as determined by germline variation of the corresponding genes) affect breast cancer susceptibility. Some studies have already investigated the role of the variation of a single pathway across the results of multiple GWAS or the variation of multiple pathways within a single GWAS in the determinism of breast carcinogenesis (80,81). However, to the best of our knowledge, this is the first time that the comprehensive collection of variants linked to breast cancer risk by means of all available GWAS (and their meta-analyses) has been employed to systematically explore the cell pathways potentially involved in breast cancer development. Our gene set enrichment analysis led to the identification of multiple pathways well known to be involved in cancer development in general (such as apoptosis, cell cycle, and signal transduction) and breast cancer in particular (such as steroid hormone pathways). As regards the latter, the estrogen receptor pathway was confirmed to play a pivotal role in the carcinogenesis of a hormone dependent neoplasm such as breast carcinoma (82), within this frame, the gene encoding the estrogen receptor alpha (ESR1) was a key influencer in the generated networks of BCPGs (see **Figure 1** and **Supplementary Table 5**). This finding might be of relevance with regard to breast cancer chemoprevention, which aims to reduce disease incidence by the administration of anti-estrogen drugs such as selective estrogen receptor modifiers (e.g., tamoxifen) (83). The selection of women who could most benefit from these risk reducing medications might be improved by genetic testing based on polymorphisms that affect breast cancer risk (84).

Another interesting piece of information yielded from data analysis is the high degree of overlap between network-guided gene set enrichment primary analysis and the pathway analysis performed with the targets of BCSGs acting as transcription factors (see **Table 1** and **Table 2**). This finding supports the hypothesis that most of the biological effect of the SNPs linked to breast cancer risk might actually be mediated by regulons governed by the transcription factors associated with those SNPs. Notably, our data confirm the results of a recent publication where investigators have identified a breast cancer risk regulatory network comprising some of the transcription factors we identified as BCPGs (85).

Besides well known cancer-related pathways (such as apoptosis, signal transduction and so on), our gene set enrichment analysis showed that germline variation of other pathways might be of particular relevance for breast cancer susceptibility, such as those involved in anti-viral immunity, degenerative diseases as well as peroxisome and ribosome activity (see **Table 1** and **Table 2**). Actually, peroxisomes are known to be linked to carcinogenesis through their production of reactive oxygen species (86), which in turn can initiate tumor development by causing DNA damage. Of special interest is also the case of genes encoding ribosome proteins, which were repeatedly enriched in our pathway and network analyses of the whole series, as well as in the analysis of pleiotropic SNPs. Indeed, it has recently been suggested that ribosome derangement may play a significant role in both development and progression of different tumor types (87,88), including breast cancer (89).

In conclusion, we present the first knowledge-base dedicated to sporadic breast cancer predisposition variants. This wealth of information can inform future studies aimed to dissect the molecular epidemiology and the molecular basis of this disease.

